# Repeated out-of-Africa expansions of *Helicobacter pylori* driven by replacement of deleterious mutations

**DOI:** 10.1101/2021.06.05.447065

**Authors:** Harry A. Thorpe, Elise Tourrette, Koji Yahara, Filipa F. Vale, Siqi Liu, Mónica Oleastro, Teresa Alarcon, TsachiTsadok Perets, Saeid Latifi-Navid, Yoshio Yamaoka, Beatriz Martinez-Gonzalez, Ioannis Karayiannis, Timokratis Karamitros, Dionyssios N. Sgouras, Wael Elamin, Ben Pascoe, Samuel K. Sheppard, Jukka Ronkainen, Pertti Aro, Lars Engstrand, Lars Agreus, Sebastian Suerbaum, Kaisa Thorell, Daniel Falush

## Abstract

*Helicobacter pylori* lives in the human stomach and has a population structure which resembles that of its host. However, *H. pylori* from Europe and the Middle East trace a substantially higher fraction of ancestry from modern African populations than the humans that carry them. Here, we used a collection of Afro-Eurasian *H. pylori* genomes to show that this African ancestry is due to at least three distinct admixture events. *H. pylori* from East Asia, which have undergone little admixture, have accumulated many more non-synonymous mutations than African strains. European and Middle Eastern bacteria have elevated African ancestry at the sites of these mutations compared to either non-segregating or synonymous sites, implying selection to remove them. We used simulations to show that demographic bottlenecks can lead to long-term segregation of deleterious mutations, despite high rates of homologous recombination, but that population fitness can be restored by migration of small numbers of bacteria from non-bottlenecked populations, leading to mosaic patterns of ancestry like that seen for *H. pylori*. We conclude that *H. pylori* have been able to spread repeatedly from Africa by outcompeting strains that carried deleterious mutations accumulated during the original out-of-Africa bottleneck.

## Introduction

*Helicobacter pylori* is the dominant bacterial member of the human stomach microbiota in infected individuals and is the aetiological agent in most cases of gastric cancer, gastric mucosa-associated lymphoid tissue (MALT) lymphoma, and gastroduodenal ulcer disease ^1^. *H. pylori* causes chronic, decades-long infections and is often acquired within the household, limiting the rate of its diffusion through human populations in comparison with more readily transmissible pathogens ^2^. Genetic variation in *H. pylori* genome sequences shows a phylogeographic pattern similar to that of its host, consistent with an inference that human and bacterial genes are often spread by the same migrations^3-6^. However, the *H. pylori* population found in Europe and other parts of Eurasia is admixed, with many strains having more than half of their DNA attributable to populations closely related to those prevalent in Africa ^3,7-9^. There is evidence for several recent human migrations out of Africa^10^, but together they have only contributed a small fraction of the ancestry of non-Africans. This discrepancy in ancestry proportions between the bacteria and their hosts implies that African *H. pylori* has been spread to Eurasia by movements of people that have left weaker signals in human DNA.

To understand why African *H. pylori* have contributed extensive ancestry within parts of Eurasia, we have assembled a collection of strains from Europe and the Middle East, from putative source populations in Africa, as well as from less-admixed strains in Asia. We infer a recent demographic history of the European and Middle Eastern strains that includes genetic drift, migration, and admixture from external sources. We show that there have been at least three admixture events from African source populations that have each contributed substantial ancestry. By examining the distribution of non-synonymous mutations in different populations, we conclude that there was a large and lasting increase in the frequency of segregating deleterious mutations during the out-of-Africa bottleneck associated with the initial spread of modern humans from Africa. When African *H. pylori* strains reached Eurasia due to later contact between humans, they, and the DNA they carried, had a fitness advantage and were able to spread. In the process, they reduced the mutational load in the newly admixed populations.

### Repeated African Admixture into Europe and the Middle East

Based on fineSTRUCTURE clustering (Figure S1), we grouped European and Middle Eastern strains into four subpopulations named hspEuropeNEurope, hspEuropeCEurope, hspEuropeSWEurope and hspEuropeMiddleEast according to the locations they were most isolated from. The first Europe in the name indicates they are subpopulations of hpEurope but for brevity we omit this part of the name in the rest of the manuscript. We investigated their sources of external ancestry by performing *in silico* chromosome painting (Figure 1A) using donor strains from three African populations (hspENEAfrica, hspCNEAfrica, hspAfrica1WAfrica) and two Asian populations (hpAsia2 and hspEAsia). Representative strains from these populations were selected from a larger collection on the basis that they showed little sign of recent admixture from other continents based on D-statistics (Table S5,S6) and as confirmed in a PCA plot (Figure S2).

**Figure 1.**
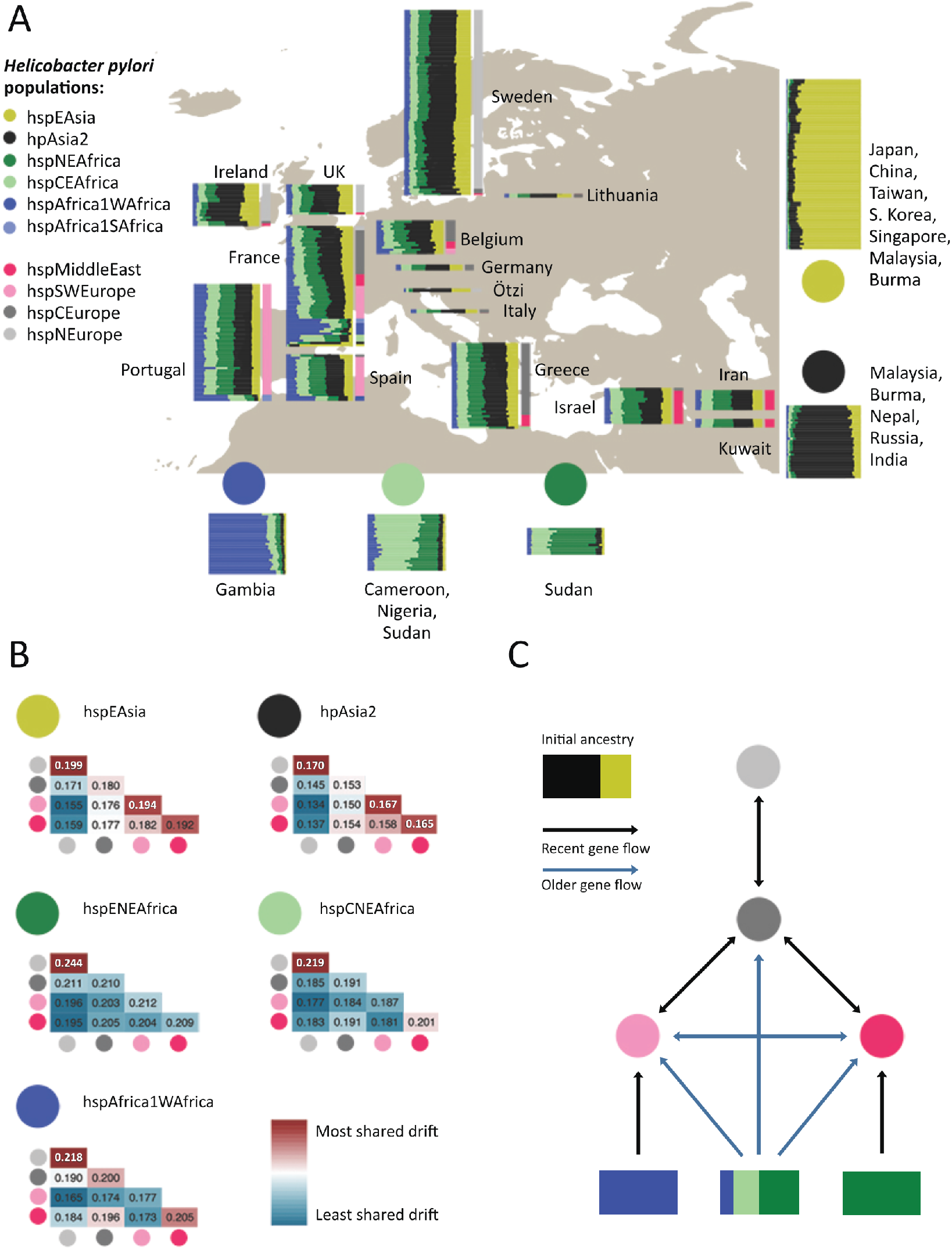
Ancestry and migration history of hpEurope isolates. (A) Painting profiles of hpEurope isolates and their putative ancestral populations from Africa and Asia showing proportion of each genome (horizontal bar) painted by each of five ancestral donor populations (circles). hpEurope isolates are grouped by country of isolation, with bars to the right indicating the H. pylori population each strain is assigned to. The representative isolates from each donor population are grouped by population, with countries of isolation listed for each group. (B) Genetic drift profiles for hpEurope subpopulations, shown separately for each ancestry component. (C) Schematic summarizing the migration and admixture history of the hpEurope subpopulations.

The chromosome painting analysis supported previous findings that there is a North-South cline in the overall proportion of African ancestry and that hpAsia2 is a closer relative of the pre-admixture population than hspEAsia^3,8^. However, all isolates are painted with a substantial and largely consistent fraction of hspEAsia (with the hpAsia2:hspEAsia ratio varying from 1:1.74 for hspNEurope to 1:2.02 for hspMiddleEast), implying that hpAsia2 is not a close surrogate for the pre-admixture population.

To investigate the demographic history of admixture further, we measured genetic drift profiles separately for each ancestry component (Figure 1B). Specifically, we compared the painting profile of pairs of admixed individuals to identify regions of the genome in which they were painted by the same donor population. For these genomic regions, we recorded if they were painted by the same specific donor strain (Methods). High rates of painting by the same donor strain indicates shared genetic drift within that ancestry component. These values are similar in interpretation to F3 values^11^, but are specific to individual ancestry components, rather than averaged across the entire genome.

Genetic drift profiles for the hpAsia2 and hspEAsia ancestry components showed a similar pattern across the four hpEurope subpopulations, as indicated by near identical pattern of colours for these two ancestry components in Figure 1B. The indistinguishable drift profiles provide evidence that within each hpEurope subpopulation, both components have been affected similarly by genetic drift. The simplest explanation is that these components are both being used to paint a single ancestry source that persisted in western Eurasia since the out-of-Africa bottleneck. Therefore, the data does not provide evidence for either ancient or more modern genetic contributions from the East.

In addition to the North-South cline, there is also an East-West ancestry cline in the source of African admixture (Figure 1A), with distinct drift patterns for each African component in the four hpEurope subpopulations (Figure 1B). Strains from hspSWEurope have the highest fractions of hspAfrica1WAfrica and this ancestry component shows low levels of drift, implying that this subpopulation has undergone recent admixture from strains closely related to those currently found in West Africa. Furthermore, there are strains from Spain, Portugal and France assigned to hspAfrica1 subpopulations and a strain in Portugal with an intermediate ancestry profile suggesting that these isolates have arrived within the last few human generations. Strains from the hspMiddleEast subpopulation have the highest fraction from hspENEAfrica and the lowest levels of drift in this component. These two populations have therefore received genetic material from Africa after the initial gene flow that has introduced ancestry across the continent. These patterns imply that there have been at least three separate admixture events involving distinct populations of African bacteria (Figure 1C). Confirmation of these results is provided by an admixture graph estimated by Treemix ^12^ (Figure S3). Treemix estimated nine migration events across the 11 populations in this study, three of which were from Africa into the European sub-populations. These three events corresponded closely to the three events found using chromosome painting.

The drift components also provided evidence about local migration. hspMiddleEast and hspSWEurope have high levels of shared drift in both Asian components, implying that prior to admixture, these two populations were closely related. hspNEurope has high subpopulation-specific drift in all ancestry components, showing that it has undergone recent genetic drift, while hspCEurope has almost none in any component, suggesting that it has been a hub for migration between populations.

The genome of a strain colonizing the Tirolean iceman, Ötzi, has been inferred to be a nearly pure representative of the pre-admixture population based on more limited data, which was interpreted as evidence that most of the admixture took place in the 5,300 years since his death ^8^. In our fineSTRUCTURE analysis, the Ötzi genome clusters with hspNEurope isolates from Ireland and Sweden with the lowest African ancestry, one of which has the same non-African ancestry proportion as the Ötzi strain in the chromosome painting. These results show that the Ötzi genome, in fact, falls within modern variation in ancestry proportions in Europe. We interpret this as evidence that substantial African ancestry had already been introduced into Europe when Ötzi lived but that ancestry proportions in particular locations have changed substantially in subsequent millennia.

Overall, the chromosome painting results show that, in addition to contemporary migrations that have introduced *H. pylori* with atypical profiles into countries such as Ireland, Portugal, France and Spain (Figure 1A), *H. pylori* have spread out of Africa at least three times (Figure 1C). Each of these migrations is sufficiently old that the DNA has been absorbed into the local gene pools, leading to a high degree of uniformity in ancestry profiles for most isolates in individual locations (Figure 1A). At least one of the early admixture events was shared between the four subpopulations, spanning Europe and the Middle East, and left traces in Ötzi’s *H. pylori* genome, while later ones, labelled as “recent” in Figure 1C, had foci in South Europe and the Middle East, respectively. Gene flow between the regional subpopulations has affected all ancestry components but has not been sufficient to homogenize ancestry proportions across the continent.

### Evidence for a role of deleterious mutations in the repeated expansion of bacteria from Africa

The ability of *H. pylori* of African origin to spread effectively in non-African populations on multiple independent occasions is unexpected, since the resident bacteria will have had an opportunity to adapt to local conditions. One potential explanation is that deleterious mutations accumulated in the genomes of strains carried by the early waves of modern humans that spread from Africa. Demographic bottlenecks associated with these migrations have been sufficient to leave an imprint on neutral genetic variation within the human genome, which indicate a reduction in effective population size in the ancestry of non-African humans around 50,000 years ago, followed by more recent expansion^13^. *H. pylori* populations also show evidence of low ancestral population sizes in East-Asian and native American populations, followed by population size recoveries^6^ consistent with strong genetic drift during the out-of-Africa and subsequent bottlenecks.

Population genetic theory ^14^ and evidence from experimental systems ^15^ has shown that demographic bottlenecks can lead to reduction in average fitness through processes such as fixation of deleterious mutations of small effect or the stochastic loss of the fittest genomes (Muller’s ratchet ^16,17^). However, although the out-of-Africa bottlenecks have had measurable impact on the pattern of segregating mutations within human populations ^18^, there is less evidence that individuals from non-African populations have accumulated a larger burden of non-synonymous mutations when measured relative to an outgroup^19^. *H. pylori* shows much higher rates of genetic differentiation between geographical regions ^4^ than its host ^13^, which most likely reflects transmission bottlenecks during spread from person to person as well as the expansion of fit clones. Consequently, demographic bottlenecks that have had modest fitness consequences for humans can potentially have more substantial effects on the bacteria they carry.

Bacteria from the hspEAsia and hpAsia2 populations, which have been through the out-of-Africa bottleneck, had a higher number of deleterious mutations segregating between strains (Figure 2A) and a higher dN/dS to the *H. acinonychis* outgroup (Figure 2B, S4B) than do the three African populations with comparable synonymous divergence levels dS. dN/dS for the three African populations ranged between 0.137 and 0.138, with Asian populations having values of 0.140 and 0.141. This corresponded to an average of 574 extra non-synonymous mutations in the Asian populations or 1 mutation for every 3 genes. dN/dS values within populations varied between 0.123-0.133 for African populations and 0.163-0.169 for the Asian ones. This difference corresponded to an average of 1216 non-synonymous differences between pairs of strains or about 2/3 of a SNP per gene. These results show that the larger number of differences in dN/dS to the outgroup in the Asian populations reflects a larger number of segregating non-synonymous variants and thus cannot simply be attributed to fixation of mutations during the out-of-Africa bottleneck.

**Figure 2.**
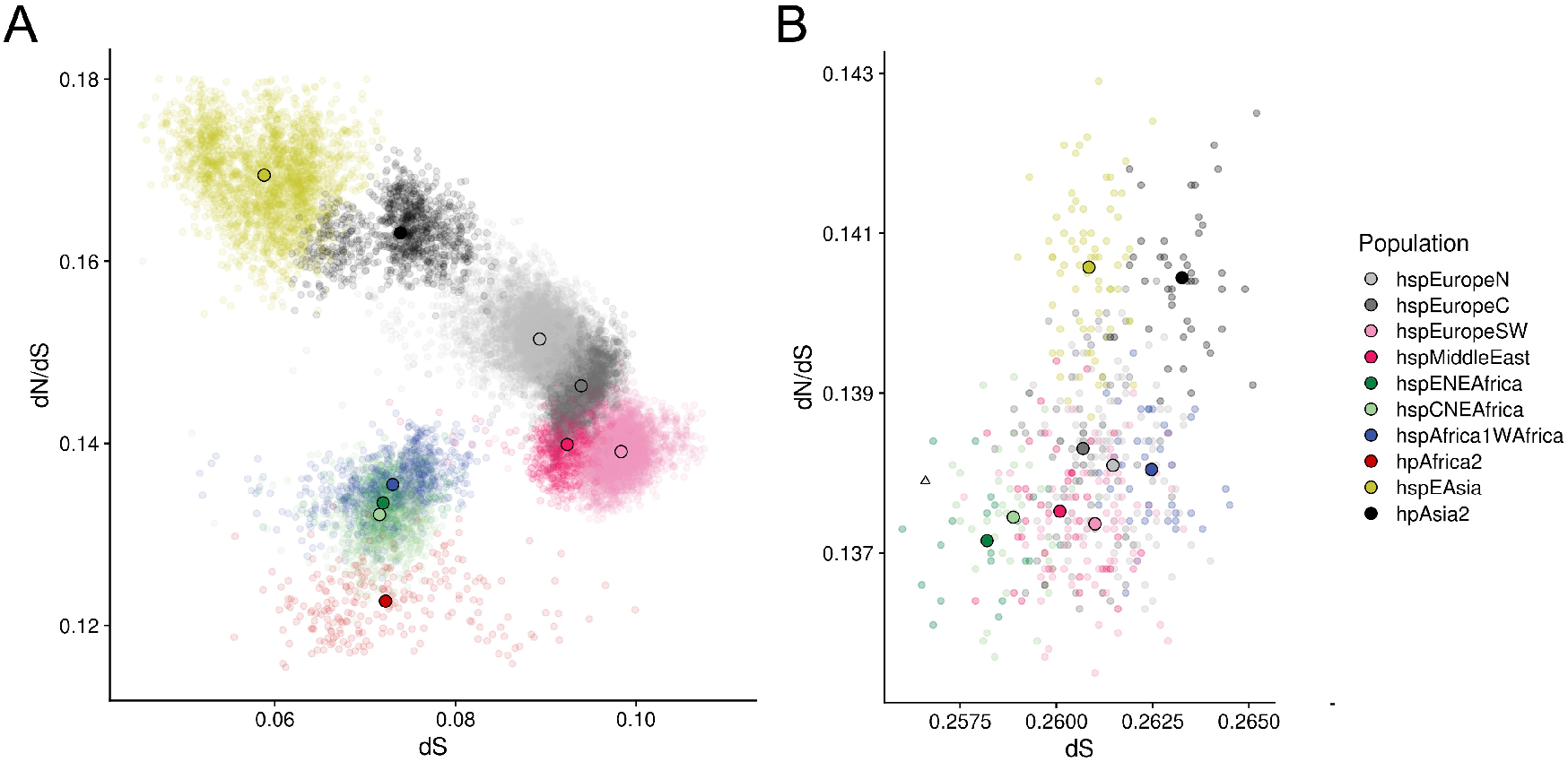
Within- and between-population divergence. (A) Within population dN/dS (y axis) plotted against dS (x axis). Small dots show pairwise distances; larger solid dots indicate population means. (B) dN/dS, calculated to the H. acinonychis outgroup, plotted against dS for isolates (semi opaque points) and populations (solid points), excluding hpAfrica2 isolates. The triangle indicates the genome from Ötzi.

We confirmed the robustness of these results in two ways. Firstly, we used GRAPES ^20^ to estimate the rate of non-adaptive non-synonymous substitutions based on the pattern of segregating mutations within each population. This approach can be applied with a folded-mutation spectrum, obviating the need for an outgroup. The results were highly concordant with those obtained for dN/dS values (Figure S5). Secondly, when an hpAfrica2 strain was used as an outgroup instead of *H. acinonychis*, similar results were obtained for dN/dS, but the range of dS values between populations was twice as large (Figure S6). This suggested that there have been ancient admixture events involving Africa2-like lineages and other African populations, while confirming the higher mutation load in Asian populations.

We also observed a reduction of the mutational load by admixture and selection within hpEurope subpopulations. Bacteria from hpEurope subpopulations had dN/dS values with the *H. acinonychis* outgroup that were intermediate between African and Asian populations but were lower (0.137-0.138) than would be predicted if they were random mixtures of African and Asian genomes with the proportions estimated by chromosome painting in Figure 1A (0.139-0.140). However, this mutation deficit is not on its own compelling evidence for a direct benefit of admixture, since it might instead reflect differences in dN/dS between the population that existed in Europe prior to admixture and the Asian populations that were used as surrogates for this ancestry in this analysis.

More specific evidence that admixture has reduced the burden of deleterious non-synonymous mutations was provided by tabulating the effect of mutations that accumulated in African and Asian populations on the ancestry of admixed bacteria in Europe and the Middle East (Figure 3). For each position in the alignment, we calculated a mutation score, which is the difference between African and Asian strains in the proportion of nucleotides that differed from *H. acinonychis* (with equal weight given to each subpopulation, see methods). We then investigated whether there is variation in overall Asian ancestry proportion (hpAsia2+hspEAsia in the chromosome painting) associated with this score. Most sites were non-polymorphic and had a mutation score of 0.0, which was therefore used as a baseline and compared independently to positive and negative scoring sites. To allow for correlations between adjacent sites, statistical significance of the regression was assessed using a gene-by-gene jackknife (methods).

**Figure 3.**
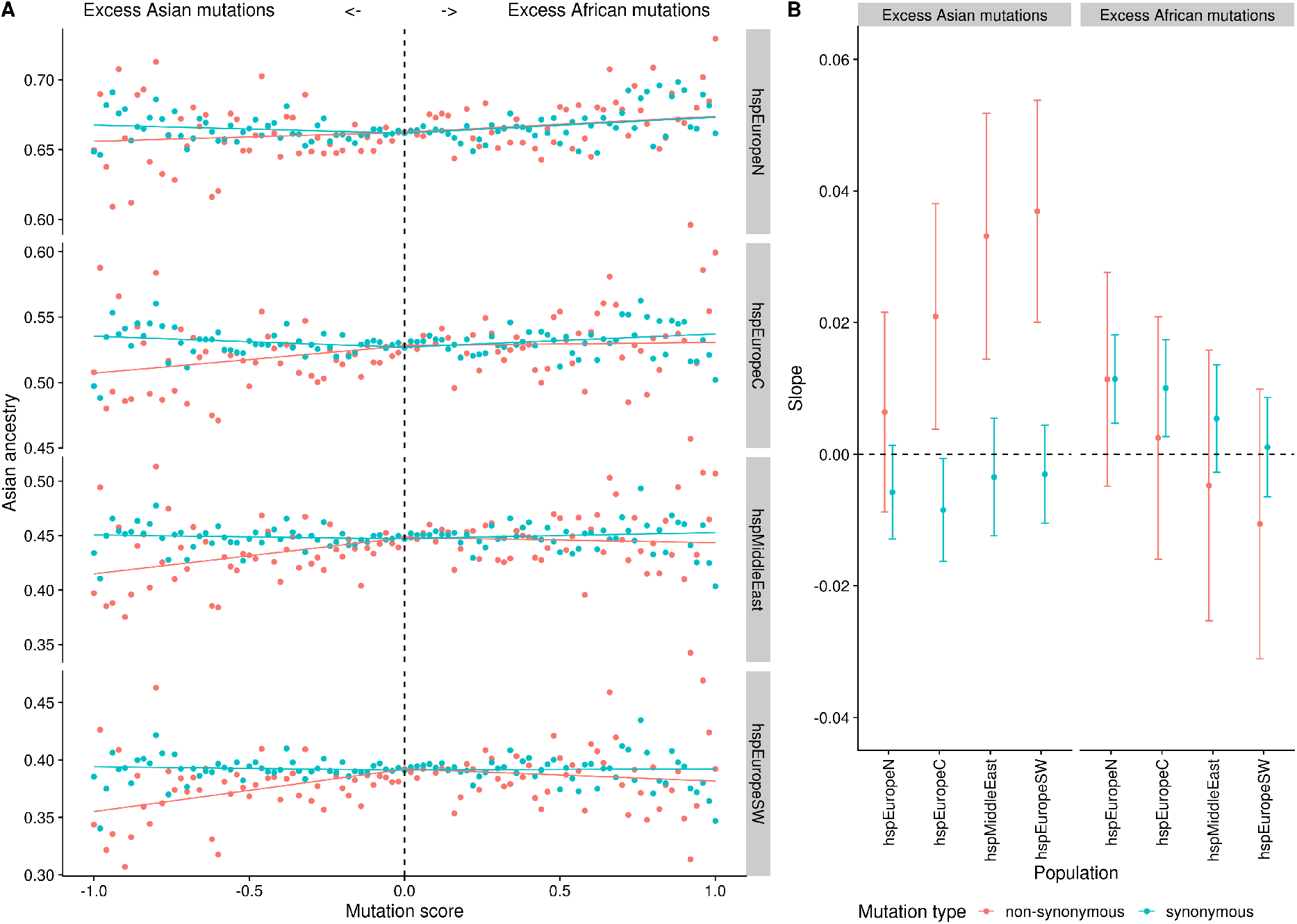
Genetic ancestry of hpEurope subpopulations as a function of mutation score. (A) Average ancestry in chromosome painting analyses plotted against mutation score (mutation frequency in African populations minus mutation frequency in Asian populations) in bins of 0.02. Regression lines were calculated separately for positive and negative mutation scores. (B) Regression slopes with 95% confidence intervals estimated using a gene-by-gene jackknife for excess Asian and excess African mutations, respectively.

*H. pylori* shows little evidence for codon usage bias^21^, so most synonymous mutations should be approximately neutral and can therefore be used as a control. A negative mutation score at non-synonymous sites is associated with a deficit in Asian ancestry, either in comparison with non-polymorphic sites or with synonymous sites with the same scores. The strongest regression slopes were estimated for the two Southern populations, and the weakest estimated for hspNEurope (Figure 3A,B). By contrast, excess mutations in African strains did not have a detectable effect on ancestry in any of the four hpEurope subpopulations, with regression slopes statistically indistinguishable from those found for synonymous positions. Thus, admixed bacteria have avoided non-synonymous mutations that accumulated in populations that have been through the out-of-Africa bottleneck, but not those that have not. We inferred that a proportion of these non-synonymous mutations are deleterious and have been selected against in the admixed population. Similar results were obtained, albeit with lower statistical confidence, when an hpAfrica2 strain was used as an outgroup instead of *H. achinonychis* (Figure S7A,B).

Deleterious mutations occurred throughout the genome and were all subject to genetic drift. A plot of mutation score versus ancestry at a gene-by-gene level showed considerable scatter (Figure S8A, Table S7), with similar results obtained when the analysis was performed for 10kb regions (Figure S8B), suggesting that the signal that we observed for mutation replacement cannot be attributed to a small number of genes. Nevertheless, it is possible that other selective forces, for example related to local adaptation, could explain some of the variation in ancestry proportion between genes.

We investigated whether specific genes were enriched for African ancestry. There was substantial variation amongst genes in the average African ancestry proportion, with strong correlations between proportions in the four hpEurope subpopulations (Figure S9), which is consistent with much of the ancestry resulting from a single shared admixture event. However, we observed no significant differences between COG categories in average ancestry proportion (Figure S10).

Genes with extreme values of average ancestry proportion were involved in diverse, often central, cellular processes. For example, low African ancestry (top ten in Table S7, each with <27% average African ancestry) genes included, *e*.*g*., those in central energy generation (HP0145, component of the Cbb3-type terminal oxidase), stress tolerance (HP0278, *ppx* exo-polyphosphatase; HP0600, *spaB* multidrug resistance), cell envelope biogenesis (HP0867, lipid A), and translation (HP1147, ribosomal proteins L19). However, the analysis did reveal overlap of low-admixture genes with genes that have highly differentiated SNPs within East Asia ^22^, specifically HP0284 (*mscS-1*), encoding a mechanosensitive channel related to osmo-tolerance and HP0250 (*oppD*), encoding an oligopeptide permease. Since the same genes are often differentiated in different continents ^22^, this overlap suggests that these genes may have been resistant to admixture due to local adaptation within Europe and the Middle East.

High-average African ancestry genes (>69%) likewise included those in central biological processes and stress tolerance, such as those of the Czc cation efflux system (HP0969-HP0970, metal ion tolerance and nickel homeostasis), purine salvage (HP0267, cytosine/adenine deaminase), and glycolysis (HP1166, glucose-6-phosphate isomerase & HP0154, enolase).

Most were not directly related to host interactions, although some with low average African ancestry might affect urease activity (HP1129, *exbD &* HP0969-HP0970, *czc*). Overall, this inspection supported that the driving force for admixture within the core genome was increased fitness, provided by replacement of stochastic deleterious mutations genome-wide, rather than single pathogenesis-related genes allowing adaptation to new host environments.

To test the hypothesis that selection on deleterious mutations can explain the observed patterns, we performed simulations of bacterial populations evolving with a constant input of neutral and deleterious mutations and homologous recombination of short tracts (Figure 4, Figure S11,S12). These simulations showed that at high recombination rates, demographic bottlenecks could generate long-term increases in the number of deleterious mutations segregating in the population (Figure 4A, Figure S12C,D), and in dN/dS measured relative to an outgroup (Figure 4B, Figure S12E,F). These patterns qualitatively match those seen in the data (Figure 2A,B).

**Figure 4.**
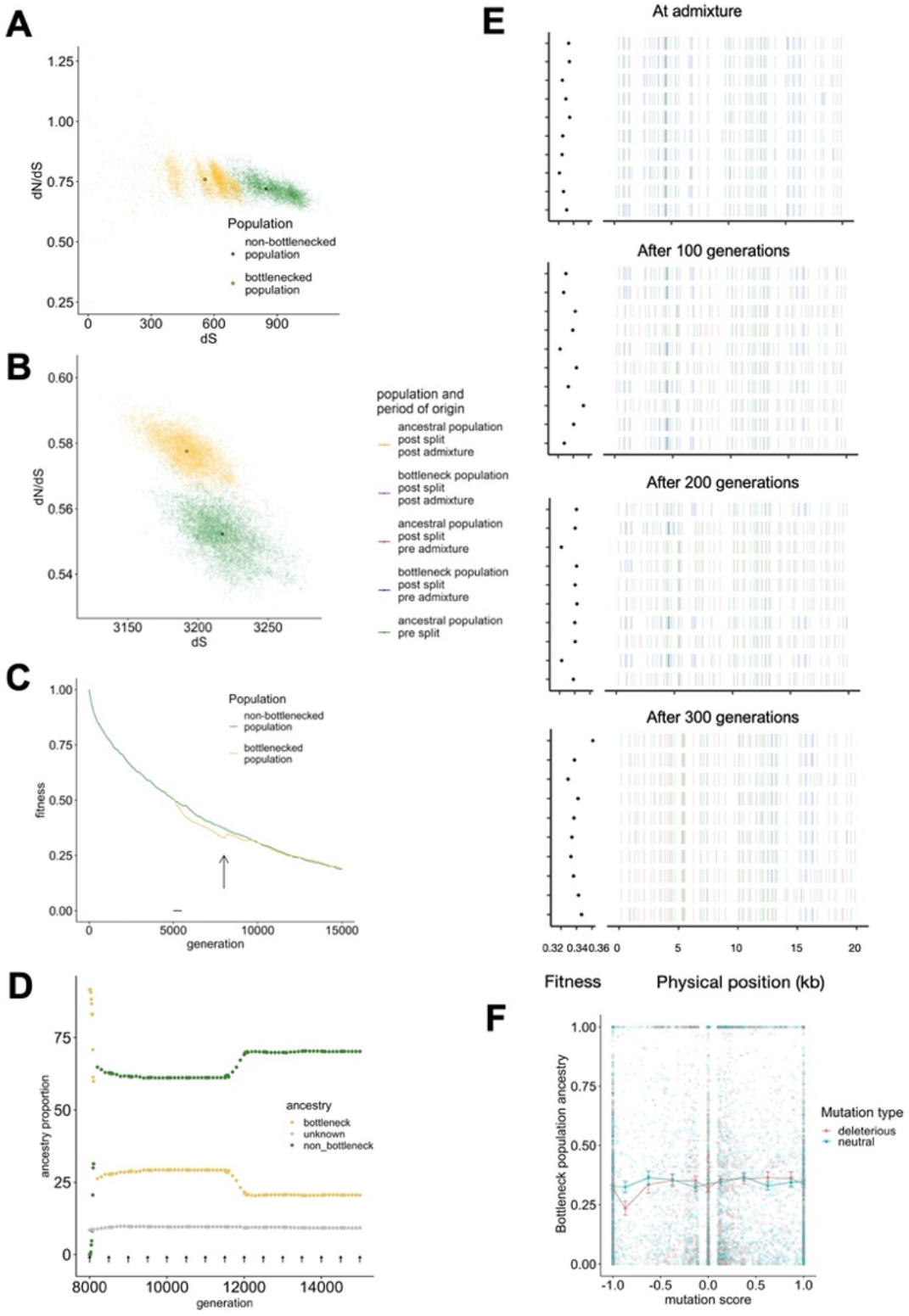
Simulations of populations under bottlenecks and admixture. (A) Within population dN/dS and (B) dN/dS calculated to the ancestor measured at generation 8000. Semi opaque points show pairwise distances; solid points indicate population means. (C) Population average fitness shown during the generations of the simulation, with bottleneck starting at generation 5000 and admixture at generation 8000, with each arrow corresponding to the migration of a single strain. (D) Proportion of bottleneck and non-bottleneck ancestry in the bottleneck population, for the generations after the beginning of admixture. Sites with unknow ancestry are shown in grey. Each arrow corresponds to the migration of one strain. (E) Ancestry painting in a sample of genomes from the bottleneck population, in the generations subsequent to admixture. Only the first 20kb of each genome is shown. (F) Average bottleneck population ancestry, at generation 8300 plotted against mutation score (frequency in the non-bottleneck population minus the frequency in the bottleneck population before the admixture begins).

In our simulations, the average fitness of the bottlenecked population underwent the largest decrease at intermediate recombination rates (Figure 4C, Figure S12A,B) and in this simulation, the bottlenecked population was susceptible to invasion from strains from the non-bottlenecked population. Once an invading strain became established in the non-bottlenecked population, the frequency of migrant DNA increased in an almost stepwise fashion leading to the generation of highly mosaic genomes (Figure 4D,E). As migrant DNA spread through the population, mutations from the bottlenecked population that were deleterious fell faster in frequency than mutations at neutral sites (Figure 4F) reproducing the dependence observed in the data (Figure 3A). This showed that the interplay between recombination and selection can explain the reduced mutational burden of the admixed populations.

Our simulations are consistent with results obtained for eukaryotic systems, which have shown that population bottlenecks can increase mutational load ^23^ and that gene flow from populations with a higher effective population size to those with a smaller one decreases the genetic load of the smaller population ^24^. This effect is particularly strong in regions of higher recombination, for which introgressed neutral/beneficial mutations will persist due to their uncoupling with the introgressed deleterious mutations. However, our simulations show that for bacterial populations the decrease of the genetic load due to admixture is only valid for intermediate recombination levels. For higher recombination rates, admixture has no effect on the fitness of the smaller population, since selection is already effective in removing deleterious mutations.

## Discussion

Deleterious mutations provide a compelling explanation for the repeated spread of African *H. pylori* into other continents, as shown by the good qualitative match between our simulation results and the pattern of diversity within contemporary *H. pylori* populations.

First, we find elevated dN/dS values in Asian populations, consistent with a higher load of deleterious mutations accumulating during the out-of-Africa bottleneck. Secondly, our simulations show that rare migrants from non-bottlenecked populations can spread and generate highly mosaic genomes, as observed in Europe and the Middle East. Third we show that ancestry from these invading lineages is higher in regions of the genome where non-synonymous mutations are segregating within the bottlenecked populations, implying that selection has acted to purge these mutations during admixture.

Deleterious mutations have been shown to be important during hybridization in several eukaryotic systems, including swordfish ^26^, trees ^27^ and Neanderthal introgression into modern humans^25^. In these systems, recombination rate variation has been shown to be crucial in determining the rate of introgression that has taken place in different regions of the genomes. To our knowledge, this is the first demonstration of a load effect in bacteria. *H. pylori* is known to recombine at an extraordinary rate ^28-30^ and this may explain why a clear signal is observable in this species.

Many of the details concerning the spread of African ancestry remain to be elucidated. The initial admixture event(s) left traces in the *H. pylori* genome of the Tirolian Iceman Ötzi, which has an ancestry profile like that found in modern day samples in Ireland and Sweden but has less African ancestry than in modern genomes from Southern Europe. Therefore, the first admixture event occurred more than 5300 years ago but the average level of African ancestry has increased substantially in the last few millennia.

*H. pylori* of different origins can mix together and recombine within extended families^31^. Over time, extensive ongoing contacts between populations within Europe would be expected to homogenize *H. pylori* ancestry proportions both within and between locations. However, we lack quantitative information on transmission dynamics that might allow us to estimate mixture dates based on the properties of the ancestry clines we observe.

An intriguing question is why some populations appear to have been resistant to invasion by DNA from African *H. pylori*. For example, within East Asia, most strains appear to come from the hpEAsia population, with very little evidence of admixture. Lack of contact with Africans does not seem a sufficient explanation, given the large number of documented contacts between East Asians and other Eurasian populations within the last several thousand years. The high burden of *H. pylori* related gastric disease in the region is notorious and hpEAsia bacteria are known for distinctive variants at virulence associated loci including *cagA* and *vacA*^32^. It is possible that strains from this population have acquired a suite of adaptations that allows them to outcompete invading bacteria despite the large mutation load within their genomes. This raises the possibility that some of the non-synonymous mutations that rose to high frequency during the bottleneck may have allowed rapid adaptation, in other words a form of evolution by shifting balance^33^.

*H. pylori* seems to be an outlier amongst bacteria in many features of its biology ^30^, including its slow rate of spread between human populations and its high mutation ^28^ and recombination rates ^29^. Our results suggest that recombination may save strains from rapid mutational meltdown but that deleterious mutations persist within populations, with the effects of bottlenecks enduring for millennia. The unusual properties of *H. pylori* make it a powerful model system for understanding how deleterious mutations interact with demographic processes and adaptive ones to mould diversity within natural populations.

## Methods

### Dataset collection

A dataset of 716 *Helicobacter pylori* whole-genome sequences was assembled, consisting of 213 newly sequenced isolates from Europe, Asia and Africa (Table S1) and selected publicly available genomes (Table S2).

### Genome sequencing

New genomes were sequenced at five different centres: Karolinska Institute, Sweden (KI), Hannover Medical School, Hannover, Germany (MHH), Hellenic Pasteur Institute, Greece (HPI), Oita University, Japan (OiU), and University of Bath, UK (UBa) (Table S1).

Genomic DNA from strains marked with KI in Table S1 was extracted using DNeasy Mini Kit (Qiagen, Hilden, Germany) following the manufacturer’s guidelines for Gram-negative bacteria. Sequencing libraries were prepared using the TruSeq Nano kit (Illumina, San Diego, CA, USA) and sequenced on the MiSeq platform, v3 chemistry, using 300 bp paired end mode.

Genomic DNA from strains marked MHH was isolated from *H. pylori* strains after 24h culture on *H. pylori* selective agar (in-house recipe) with the Genomic Tip 100/G (Qiagen, Hilden, Germany). Nextera XT libraries were generated and sequenced in three different runs on MiSeq 2×300bp paired (Illumina, San Diego, CA, USA), as recommended by the manufacturer. All quantification steps of gDNA and NGS libraries were done with Qubit dsDNA HS Assay Kit (Invitrogen, ThermoFisher Scientific, Carlsbad, CA, USA).

For strains marked with HPI, adapter-compatible DNA was prepared using Ion Xpress™ Plus Fragment Library Kit and enzymatically fragmented for 5-12 minutes, resulting in a median fragment size of 350-450 bp and the libraries were prepared using the Ion Plus Fragment Library Kit. The resulting 400 bp insert libraries were used to prepare enriched, templatepositive Ion PGM™ Hi-Q™ View Ion Sphere Particles (ISPs) with the Ion OneTouch™ 2 System. 850-flows sequencing was performed using the Ion PGM™ Hi-Q™ View Sequencing Kit with the Ion 318™ Chip Kit v2.

For genomes marked with UBa, genomic DNA was quantified using a NanoDrop spectrophotometer, as well as the Quant-iT DNA Assay Kit (Life Technologies, Paisley, UK) before sequencing. High-throughput genome sequencing was performed using a HiSeq 2500 machine (Illumina, San Diego, CA, USA).

Genomic DNA from strains marked with OiU was extracted using DNeasy Blood & Tissue kit (QIAGEN, Hilden, Germany). DNA concentration was measured using QuantusTM Fluorometer (Promega). High-throughput genome sequencing was performed either on Hiseq 2000 (2 × 100 or 2 × 150 paired-end reads) or Miseq (2 × 300 paired-end reads) sequencer (Illumina, San Diego, CA) following the manufacturer ‘s instruction.

### Primary bioinformatics analysis

For the KI genomes, the raw sequencing reads were quality trimmed and filtered using TrimGalore! (http://www.bioinformatics.babraham.ac.uk/projects/trim_galore/) applying a minimum q-score of 30, and *de novo* assembled using SPAdes ^34^ with the –careful option. Contigs with very low coverage and that were shorter than 500 bp were discarded prior to annotation.

For the MHH genomes quality filtering was done with Trimmomatic version 0.36 ^35^ and the assemblies were performed with SPAdes genome assembler v. 3.9.0 and resulted in number of all contigs from 26 up to 117. Assemblies were quality controlled with QUAST^36^.

For the HPI genomes, quality control of raw sequencing reads was performed using FASTQC (https://www.bioinformatics.babraham.ac.uk/projects/fastqc). Unbiased *de novo* assembly was performed using SPAdes genome assembler v.3.5.0in default mode.

For the UBa genomes, raw sequencing reads were quality trimmed and filtered using Trimmomatic version 0.33 and the 100 bp short read paired-end data was assembled using the *de novo* assembly algorithm Velvet version 1.2.08^37^. The VelvetOptimiser script (version 2.2.4) was run for all odd k-mer values from 21 to 99. The minimum output contig size was set to 200 bp with default settings, and the scaffolding option was disabled.

For the OiU genomes, Trimmomatic v0.35 was used to remove adapter sequences and low-quality bases from raw short reads data. Trimmed reads were then de novo assembled to produce contigs using SPAdes genome assembler v3.12.0 with the -careful option to reduce mismatches in the assembly. The minimum contig length was set to 200 bp.

Annotation of both newly sequenced draft genomes and publicly available sequences was performed using the prokka annotation pipeline v 1.12 ^38^ using the most recent version of the 26695 genome ^39^ as primary annotation source.

Genome size and contig/scaffold number was collected from the prokka annotation output using the MultiQC tool ^40^ and collected into Table S3. All newly sequenced genomes were submitted to GenBank under BioProject PRJNA479414.

All strains, their population designations and their role in the respective analyses are shown in Table S4.

### Sequence comparison and alignment

All isolates were mapped to the 26695 genome (NC000915.1) using the Snippy software version 3.2-dev (https://github.com/tseemann/snippy). The resulting core genome, which was collected with the same tool, contained 287 746 core SNPs from 979 771 variant sites.

### FineSTRUCTURE

We inferred population structure among the strains based on the genome-wide haplotype data of the reference-based alignment to 26695 described above, using chromosome painting and fineSTRUCTURE ^41^ according to a procedure of our preceding study that applied them to *H. pylori* genome^42^. Briefly, we used ChromoPainter (version 0.04) to infer chunks of DNA donated from a donor to a recipient for each recipient haplotype and summarized the results into a “co-ancestry matrix”, which contains the number of recombination-derived chunks from each donor to each recipient individual. We then ran fineSTRUCTURE (version 0.02) for 100,000 iterations of both the burn-in and Markov Chain Monte Carlo (MCMC) chain, in order to conduct clustering of individuals based on the co-ancestry matrix.

### Choice of donor and recipient strains and chromosome painting

D-statistics were calculated for strains assigned to each of the five ancestral populations hspEAsia, hpAsia2, hspCNEAfrica, hspENEAfrica and hspAfrica1WAfrica. D-statistics were calculated using popstats (https://github.com/pontussk/popstats) and specifying individual A as SouthAfrica7 (hpAfrica2), individual B as GAM260Bi (hspAfrica1WAfrica), individual Y as F227 (hspEAsia) and individual X. In this comparison, negative D-statistics imply more African ancestry in the strain designated as individual X than in F227. D-statistics values can be found in Table S5 and Table S6.

A subset of strains from each of the five ancestral populations were chosen as donors to get groups of similar size and were selected based on the fineSTRUCTURE analysis to get good representativeness over the donor populations. Some hpAsia2 strains showed signs of elevated African admixture based on negative D-statistics values and these strains were not selected as donors.

We conducted chromosome painting of 646 recipient strains (belonging to hspNEurope, hspCEurope, hspSWEurope, hspMiddleEast, hspENEAfrica, hspEAsia, hspCNEAfrica, hspAfrica1WAfrica, hpAsia2, and hpAfrica2). For this purpose, we used ChromoPainterV2 software ^41^. For each recipient population, we calculated site-by-site average copying probability from each of the five donor populations. Gene-by-gene averages were also calculated by averaging of the sites in each gene for the 790 genes in the alignment.

Regressions between gene-by-gene averages in different hpEurope subpopulation were calculated using the aq.plot() function in R.

### Shared drift estimation

We used the chromosome painting analysis to investigate shared genetic drift profiles. We calculated a separate drift profile for each pair of hpEurope subpopulations, including a within-population profile. For each profile, we calculated separate drift values for each of the five ancestry components (with the components shown in separate triangles in Figure 1B). For example, to calculate the shared drift profile of hspNEurope and hspSWEurope, we took each combination of pairs of strains from the two populations and asked whether they used donors from the same population at each site in the genome. We also tabulated whether they used exactly the same donor strain. The drift value for that pair of populations for that ancestry component is the ratio of shared donor to shared strain, summed over all pairs and sites in the genome.

### dN/dS calculations

From the fineSTRUCTURE analysis, a sub-dataset was collected consisting of the European strains assigned to the hpEurope populations, together with a representative selection of strains from the ancestral populations hpAfrica2, hspAfrica1WAfrica, hspCNEAfrica, hspENEAfrica, hpAsia2, and hspEAsia. The *H. acinonychis* genome was added to this dataset to provide an outgroup. For a detailed list of which strains that were included in these analyses, see columns in Table S2,S4. dN/dS was estimated pairwise between these strains from core genome alignments using the method of Yang and Nielsen^43^, as implemented in Paml v4.7.

To calculate the numbers of excess mutations observed in the Asian populations, the mean number of non-synonymous mutations was calculated for African and Asian populations, and the difference was then multiplied by a correction factor of 1.37 to account for the loss of some coding sites in the core genome alignment (1.11Mb) compared with the reference genome (1.52Mb).

### Ancestry and mutation score analysis

Using *H. acinonychis* as a reference, non-synonymous and synonymous mutations were called against representative strains from the African and Asian populations. For this analysis the ancestral populations were combined to reduce the ancestry components to either African (hpAfrica) or Asian (hpAsia). For each site, the frequencies of these mutations within each ancestral population were calculated to give a score between 0 and 1. These scores were then combined by subtracting the hpAsia scores from the hpAfrica scores to give a score between -1 and 1, where -1 = fixed in hpAsia and absent in hpAfrica, 0 = equal frequencies in hpAsia and hpAfrica, 1 = absent in hpAsia and fixed in hpAfrica. Thus, these scores represent the mutational load for each site in the ancestral populations. We then combined these site-by-site scores with the site-by-site chromosome painting data for the European populations so that each site had an estimate of ancestry component from both Asia and Africa, and a mutational score representing the mutational load in the ancestral populations.

We then computed linear regressions of hpAsian ancestry against mutation score for each European population, and we did this for the positive (excess African) and negative (excess Asian) mutation scores separately. To confirm that the regression slopes were not driven by a small number of outlier genes we conducted a gene-by-gene jackknife by repeating the regressions but removing all the sites from a single gene each time. From this distribution of slope values we calculated the pseudovalues as pseudovalue=slope-((n-1)*(j_slope-slope)). We then calculated the mean and confidence limits of these pseudovalues as mean(pseudovalues) ± quantile(p=0.05/2, df=n-1) * se(pseudovalues).All data manipulation and statistical analysis was performed in R 3.6.1. Packages from the ‘tidyverse’ collection were used extensively, and ggplot2 was used for plotting.

### PCA

The PCA was realized using the software PLINK v1.90 ^44^on biallelic SNPS from the core genome CDS of *H. pylori*. The filtered SNPs were first LD pruned using the same software to get independent SNPs.

### Rate of non-adaptive non-synonymous substitutions relative to neutral divergence

The rate of non-adaptive non-synonymous amino-acid substitutions relative to neutral divergence was calculated based on the method of Galtier^20^ and implemented in the software GRAPES ^20^. The software was run with folded SFS obtained from the core CDS as input and no divergence data. The synonymous and non-synonymous folded SFS were obtained using the PopGenome library ^46^ in R^45^, and the number of synonymous and non-synonymous sites was estimated based on the reference core genome sequence.

### Admixture graphs

Admixture graphs between the different populations were obtained using the software Treemix v1.12 ^12^. Treemix was run with a number of migration edges between 0 to 15, with 10 replicates for each number of edge and hpAfrica2 was set as the outgroup. The final number of migration edges was chosen as being the smallest number that allowed 99.8% of the variance to be explained, which is the same criterion used by^12^.

### Simulation of bacterial populations evolving under deleterious mutation pressure

We used SLiM v3 ^47^ to simulate the accumulation of deleterious mutations in bacterial populations. Each bacterial genome was a single haploid chromosome of length 1.6 Mb, with a mutation rate per site per generation of 5×10^−7^. This corresponds to about 1/20 of the mutation rate per year estimated from *H. pylori* data ^28 48^. Half of the mutations are neutral, with the other half being deleterious with six different selection coefficients (-5×10^−3^, -2×10^−3^, -10^−3^, -5×10^−4^, -2×10^−4^, -10^−4^), each accounting for 1/12 of the mutations. There is no back mutation and mutations have a multiplicative effect on the fitness.

At the beginning of the simulation, there is no variation in the population. These parameters resulted in an intermediate mutation load for each strain. Higher mutation rates would have led to excessive load in each bacterial generation, while smaller selection coefficients would have led to individual deleterious mutations behaving as if they were neutral.

In each generation, recombination happens between strains from the same population via the transfer of one segment from a donor strain into a recipient strain, with the length of the block taken from an exponential distribution. We used three different mean values, 0, 5,000 and 50,000bp. *H. pylori* has a high rate of import of tracts with mean around 400bp ^49^ but simulating larger tracts is more computationally efficient than simulating many tracts.

A single population of size 10,000 evolves for 5,000 generations, by which time the population was in approximate mutation-selection equilibrium. 1,000 of the strains are moved into a second population, which retains this size for 500 generations before expanding to size 10,000. This is substantially below estimated ancestral population sizes of 2,000,000 or more for *H. pylori* ^6^ but sizes substantially in excess of this are difficult to simulate due to memory and computation time issues. From generation 8,000 onwards, migration from the non-bottleneck to the bottleneck populations happens at a rate of one strain every 500 generations. The simulations are completed after 15,000 total generations. Simulation output was analysed using python and R. For each population, we calculated its average fitness, its fitness variance, its dN/dS to the ancestor and within population dN/dS. Subsequent to admixture, the ancestry proportion from each population was tabulated based on mutations that had arisen during the separate evolution of the populations prior to mixture. For sites without such mutations, their origin was assumed to be the same as the closest mutation of known ancestry, provided that the mutation was within 1kb.

Otherwise, its origin was recorded as unknown.

The SLiM, python and R scripts can be found at https://github.com/EliseTourrette/scriptPubli/tree/main/ThorpeEtAl2021.

## Acknowledgements

The authors thank Sabrina Woltemate for technical assistance. This work was supported by Sequencing Grants-in-aid for Scientific Research from the Ministry of Education, Culture, Sports, Science, and Technology (MEXT) of Japan (221S0002 and 18KK0266) to YY. FFV is financed by FCT through Assistant Researcher grant CEECIND/03023/2017. KT and the sequencing of KI isolates was supported by Erik Philip-Sörensen Foundation grant G2016-08, and Swedish Society for Medical research (SSMF). All primary bioinformatics and parts of the comparative genomics were performed on resources provided by Swedish National Infrastructure for Computing (SNIC) through Uppsala Multidisciplinary Center for Advanced Computational Science (UPPMAX) under projects snic2018-8-24 and uppstore2017270.

Work by SS was supported by the German Research Foundation (DFG, project number 158 989 968–SFB 900/A1) and by the Bavarian Ministry of Science and the Arts in the framework of the Bavarian Research Network ‘New Strategies Against Multi-Resistant Pathogens by Means of Digital Networking – bayresq.net. DF was supported by Shanghai Municipal Science and Technology Major Project No. 2019SHZDZX02.

## Supplementary Figures

**Figure S1.**
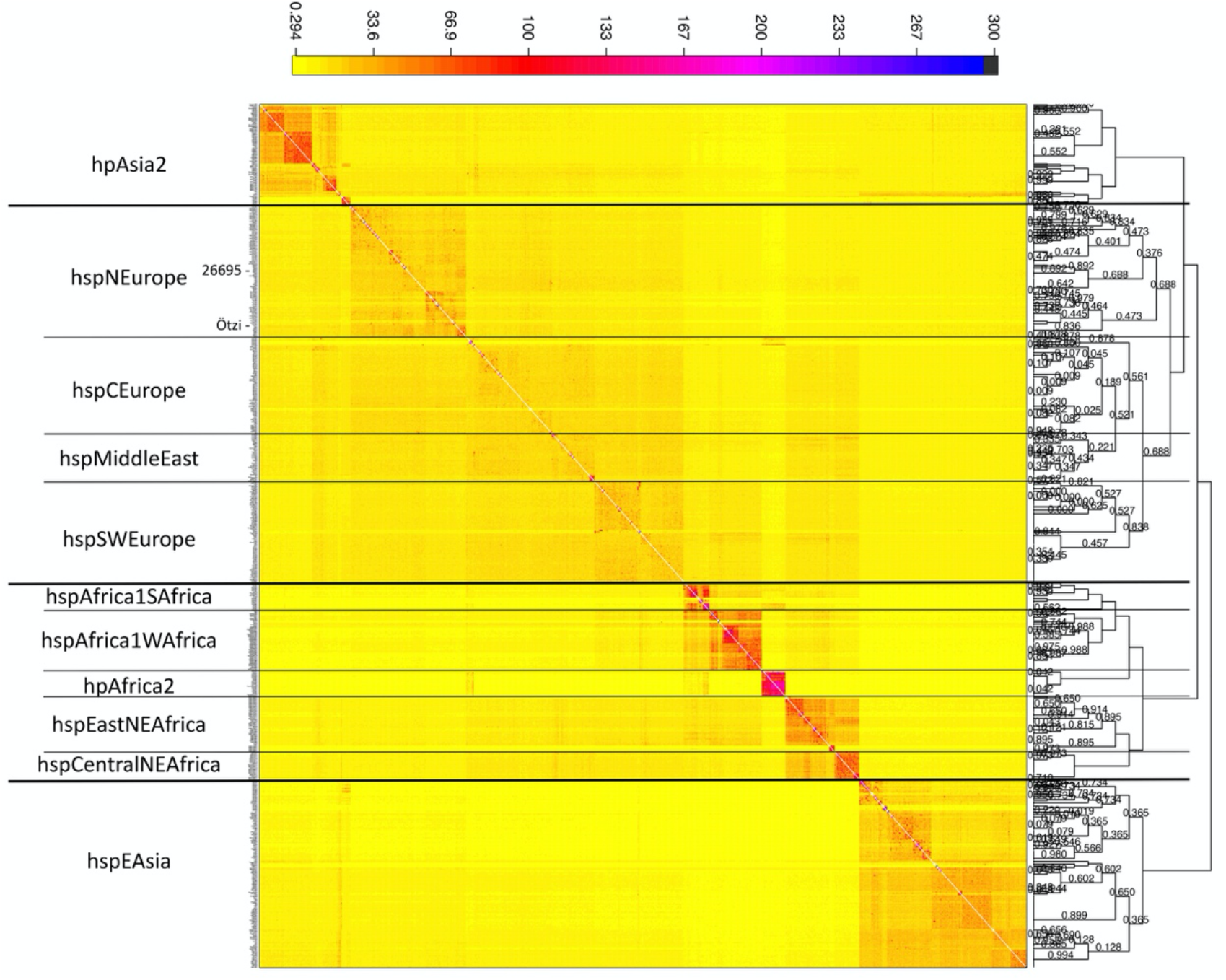
heatmap showing populations inferred by fineSTRUCTURE. For each individual, the number of chunks donated by other individuals are shown according to the colour scale at the top. Note the position of Ötzi within hpsNEurope. The tree on the right shows hierarchical clustering of fineSTRUCTURE populations.

**Figure S2:**
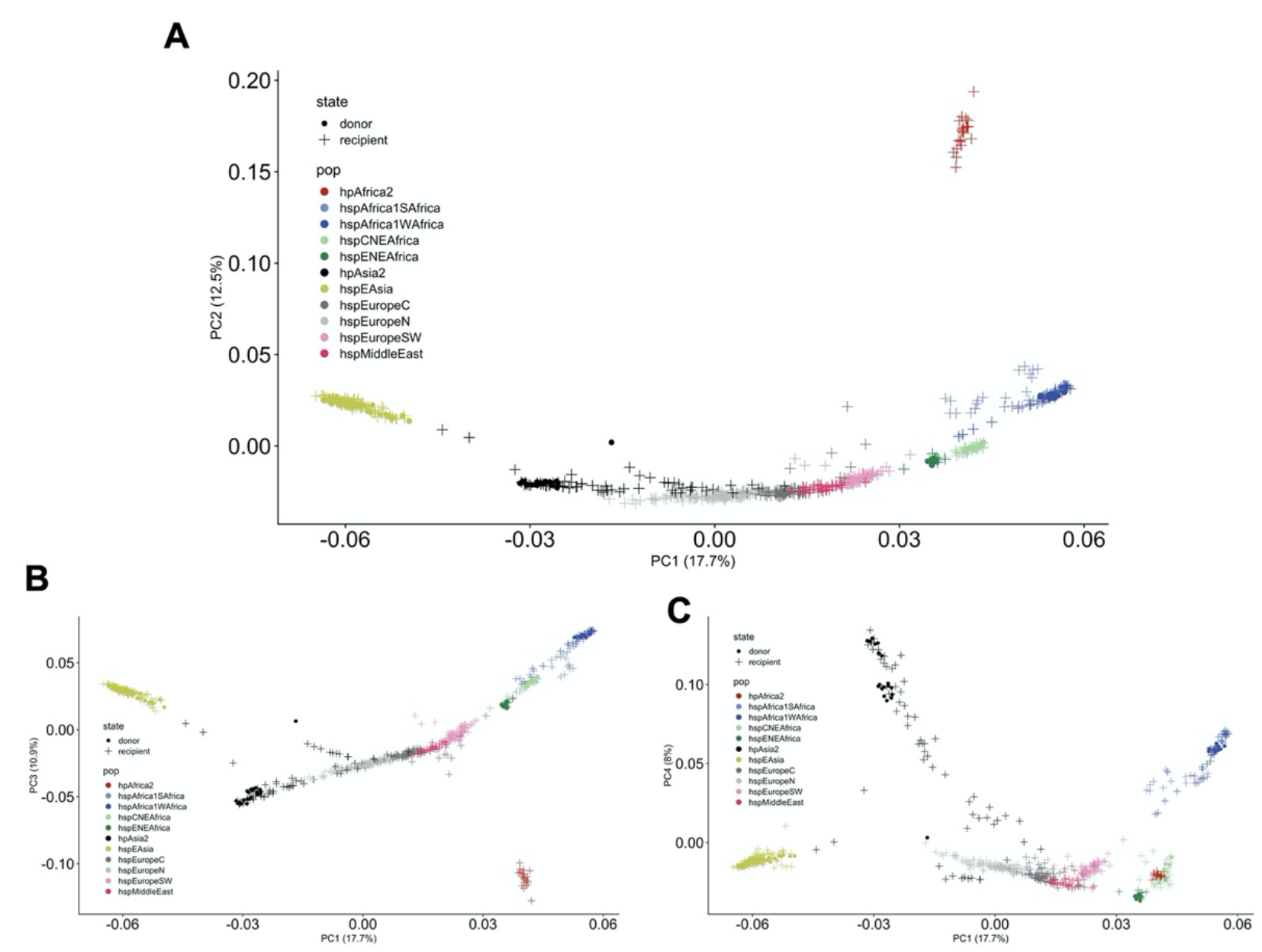
First four components of a PCA showing the different African, Asian and European populations. Strains used as donors in the fineSTRUCTURE analysis are shown as circles while the recipient are shown as crosses. (A) PC2 vs PC1, (B) PC3 vs PC1 and (C) PC4 vs PC1.

**Figure S3:**
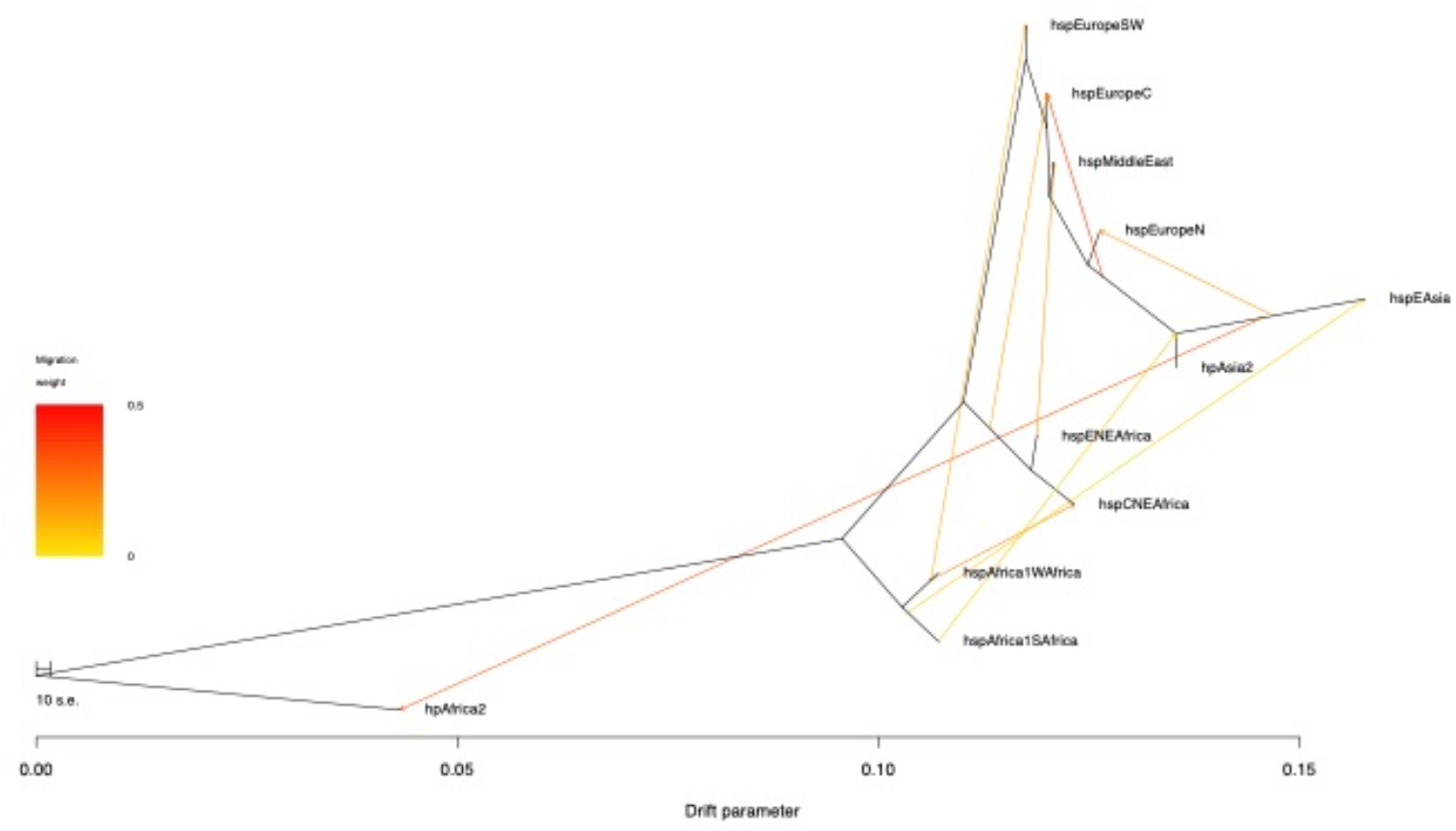
Admixture graph for the different populations, with nine migration edges – the optimum number of migration edges. The population hpAfrica2 was set as the outgroup. The arrows represent the gene flows between the different branches, their color indicating the weight of the migration edge.

**Figure S4.**
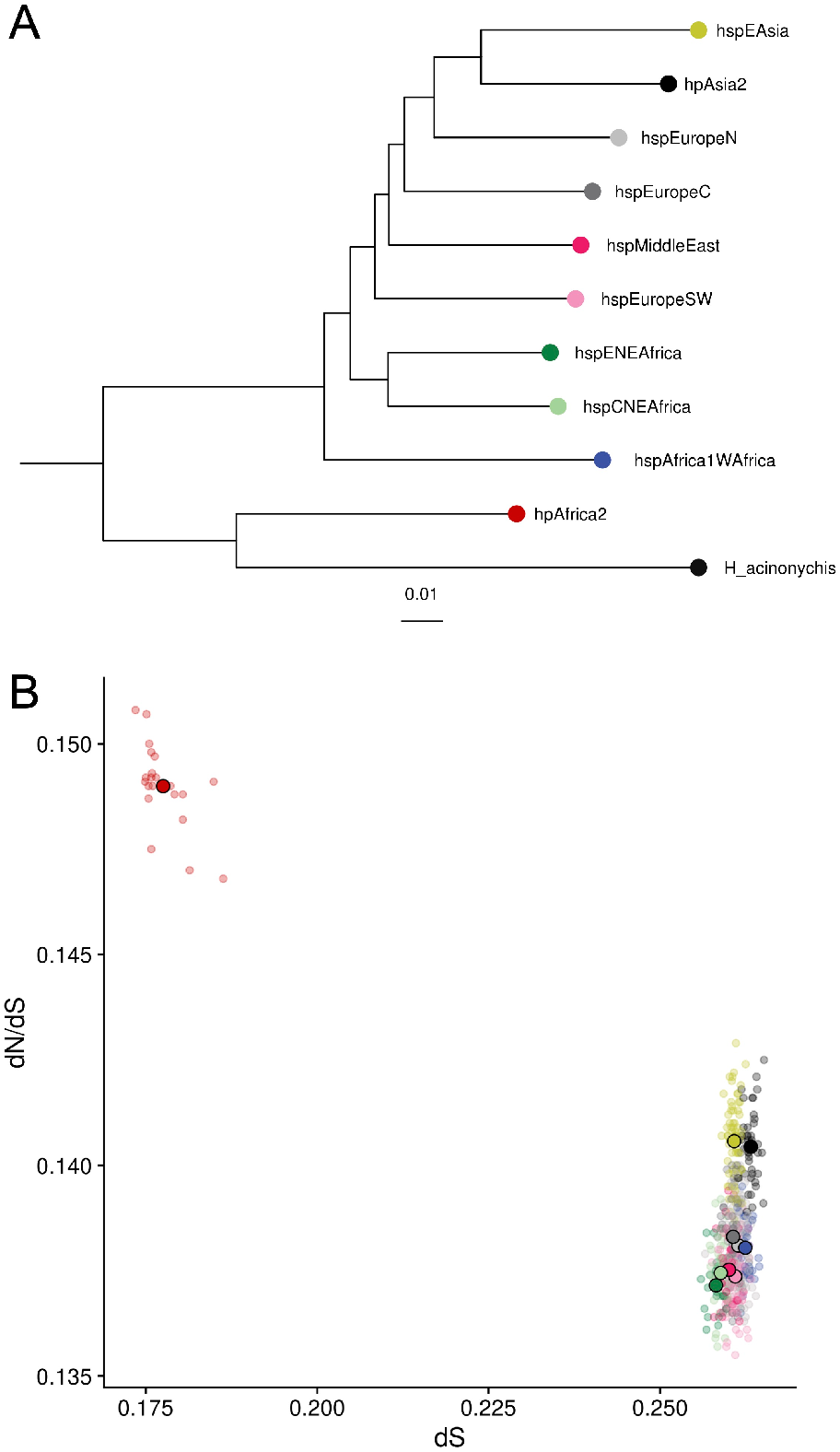
Relationships amongst populations. (A) Neighbor joining tree based on genetic distances between populations. (B) dN/dS plotted against dS to H. acinonychis outroups for all populations, including Africa2.

**Figure S5:**
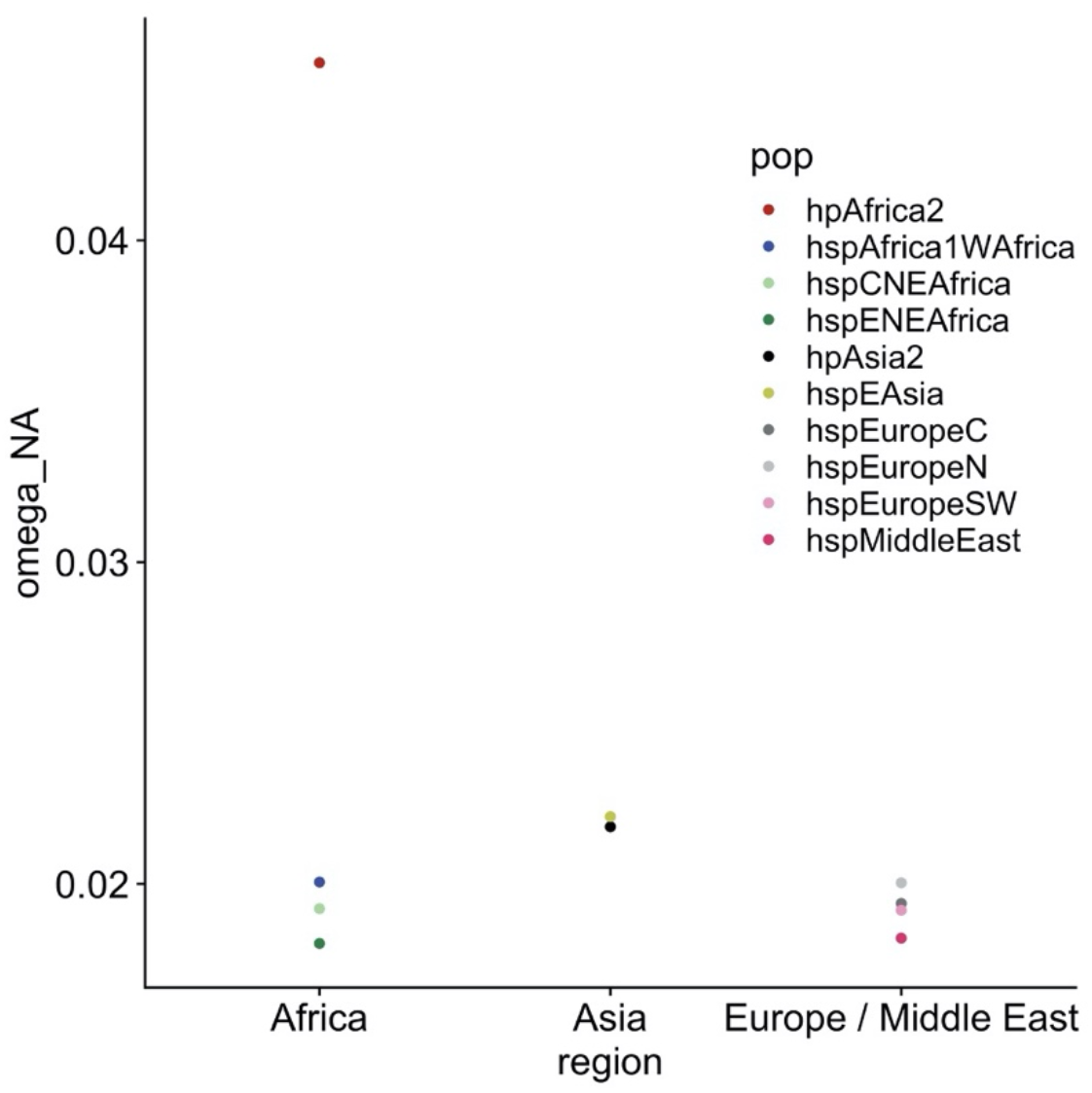
Rate of non-adaptive non-synonymous amino-acid substitutions relative to neutral divergence for the different populations, separated based on the different geographical regions on the x-axis.

**Figure S6:**
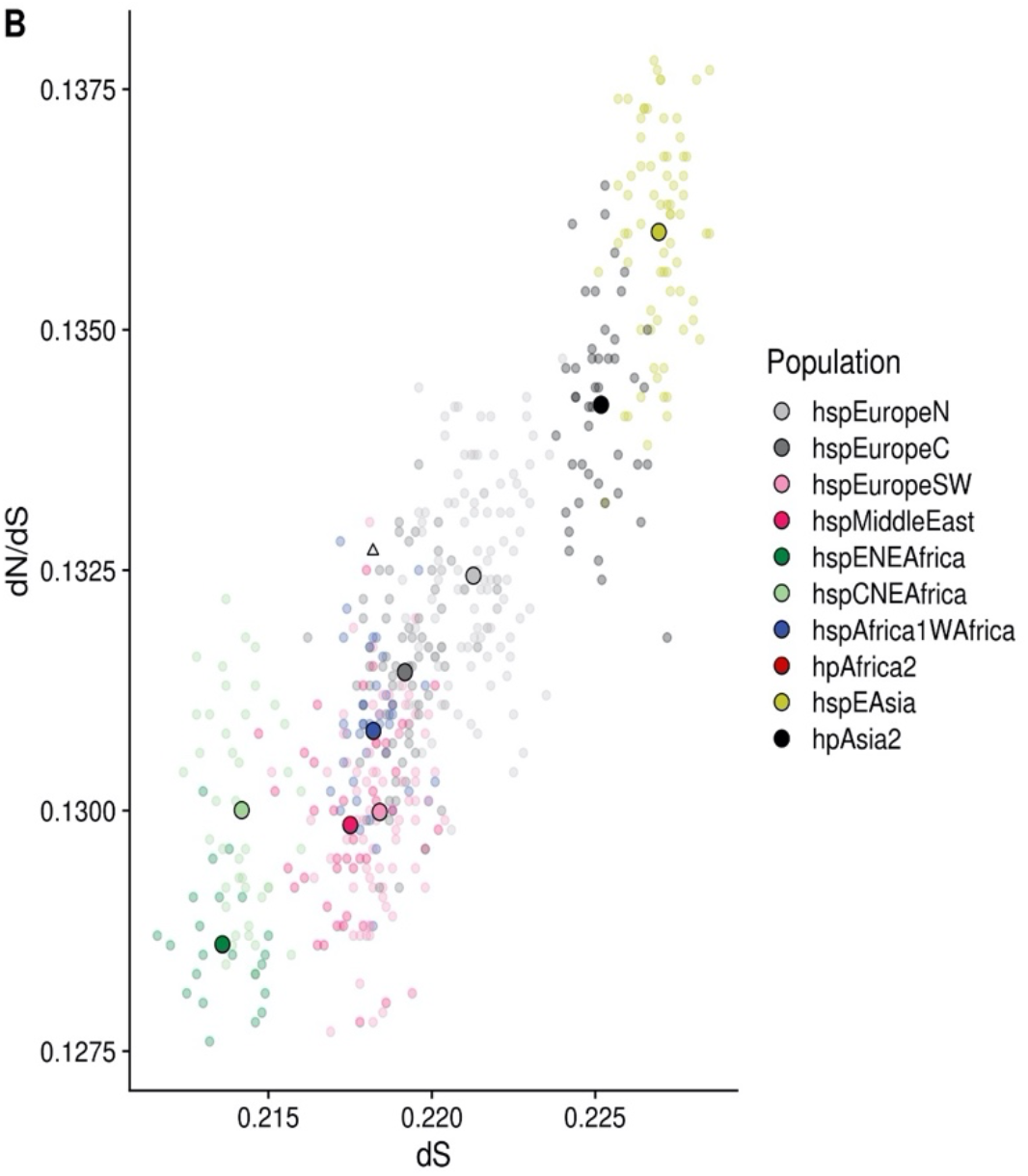
Same as Figure 2B but with hpAfrica2 set as the outgroup.

**Figure S7:**
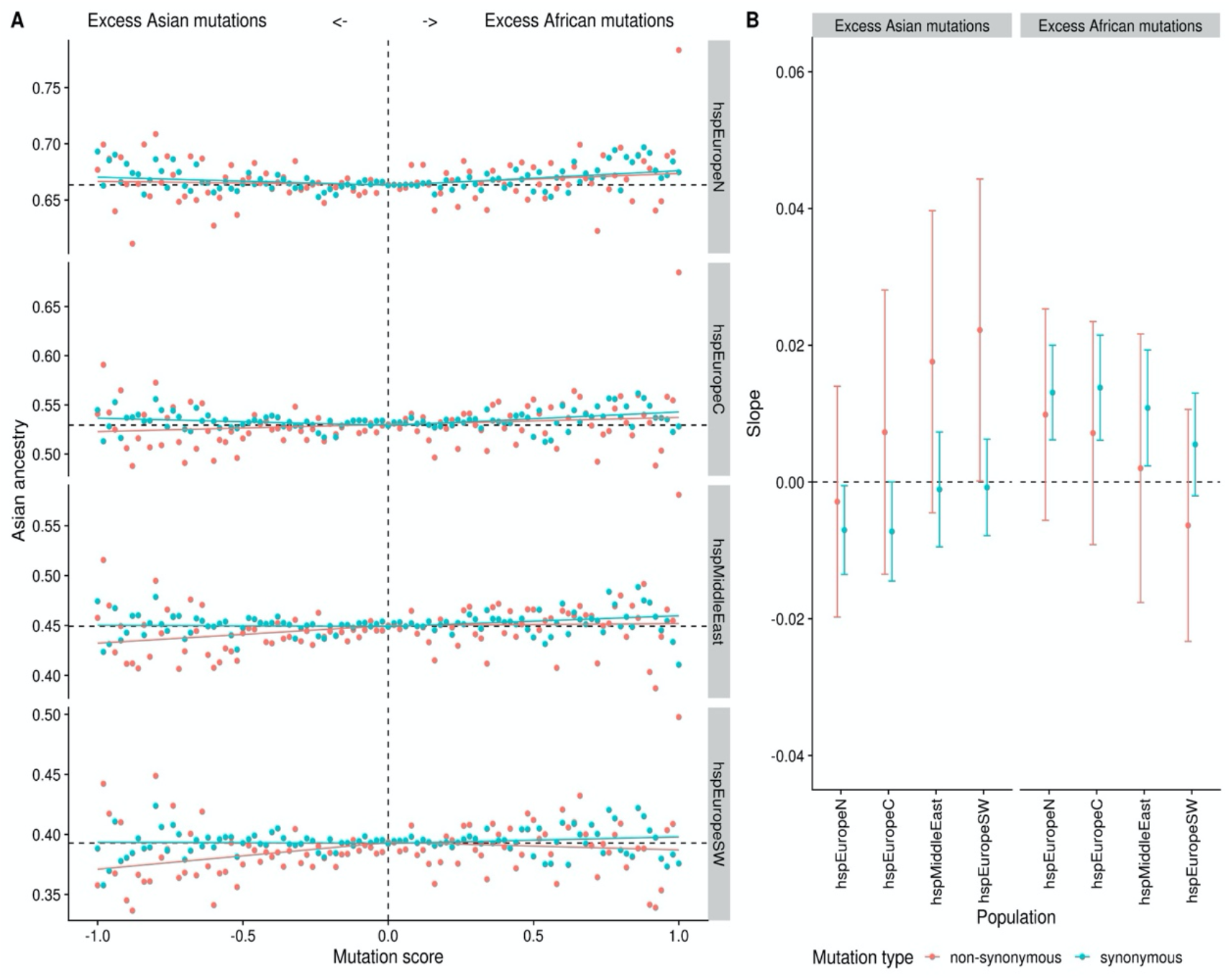
Same as Figure 3 but with hpAfrica2 set as the outgroup.

**Figure S8:**
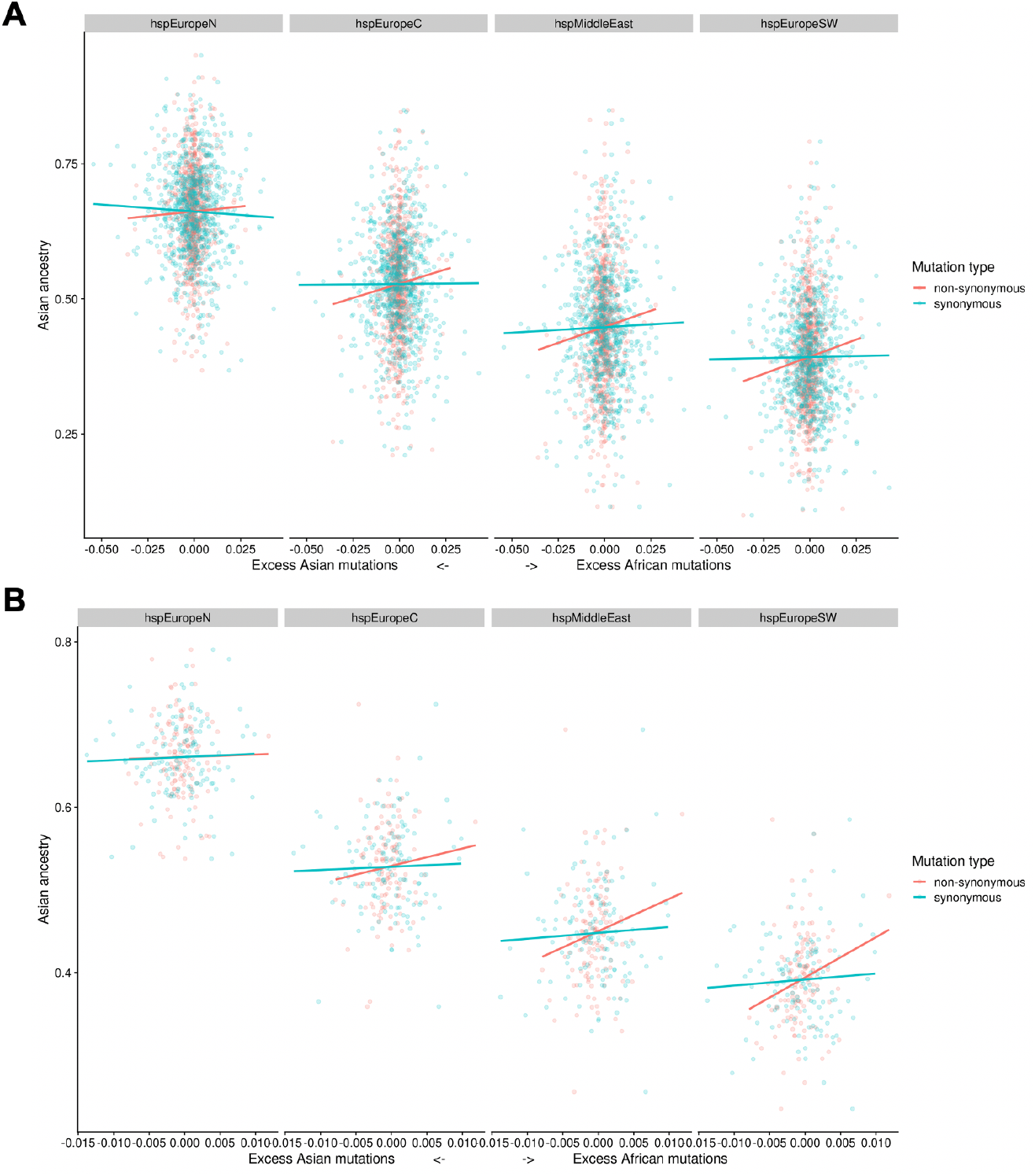
Average ancestry in chromosome painting analyses plotted against mutation score (mutation frequency in African population minus mutation frequency in Asian population) (A) by genes and (B) by 10 kb bins. In both cases, only the regression in hspEuropeSW, for the non-synonymous sites, is significative. Regression lines are calculated separately for positive and negative mutation scores

**Figure S9.**
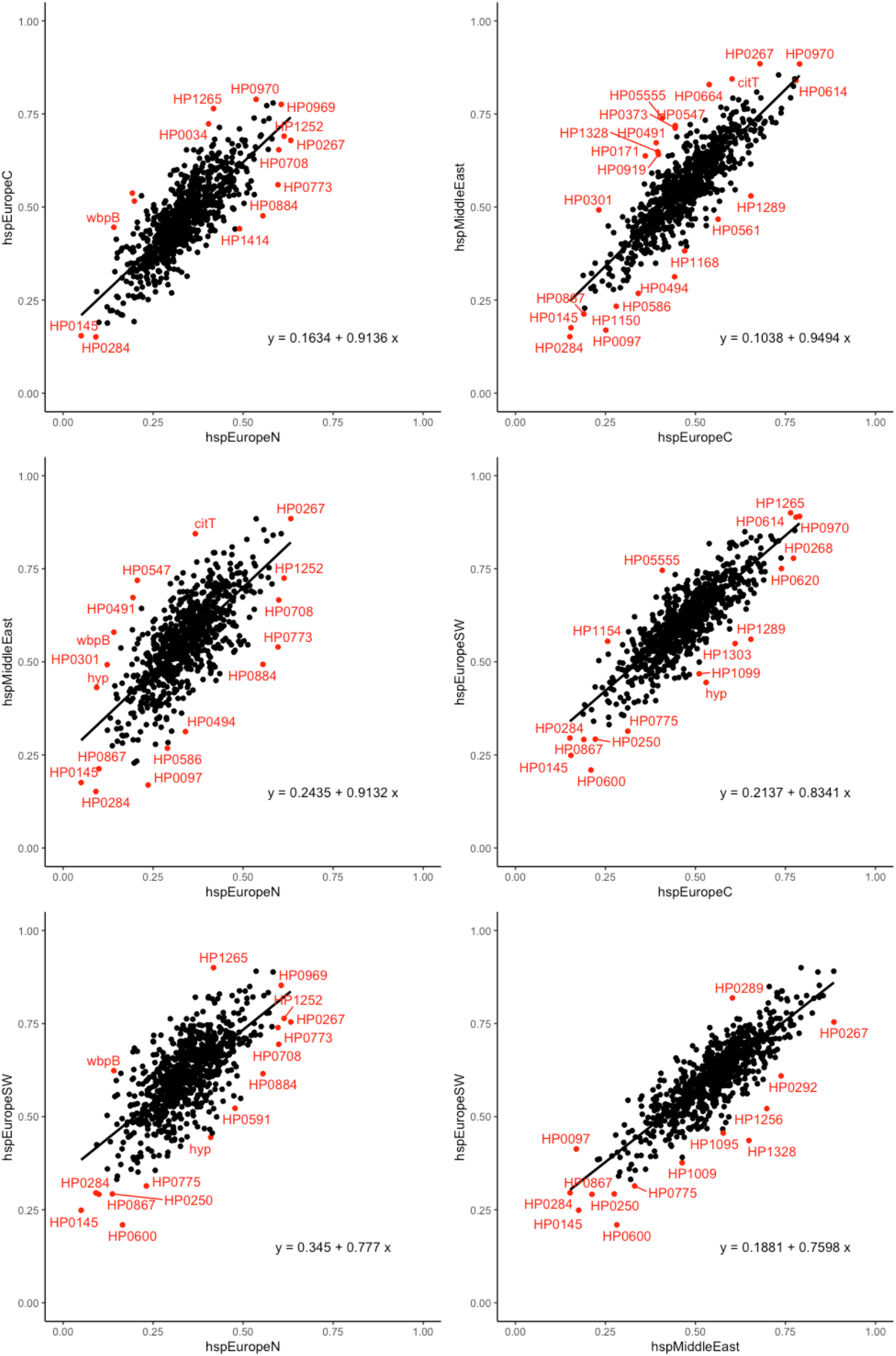
Average African ancestry proportions of genes. Each panel shows the relationship between the proportion of ancestry assigned to Africa in the chromosome painting for each gene, for pairs of hpEurope subpopulations. Outliers from the regression slope are calculated based on robust Mahalanobis distances using the R function aq.plot(data, quan =1,alpha =0.015).

**Figure S10.**
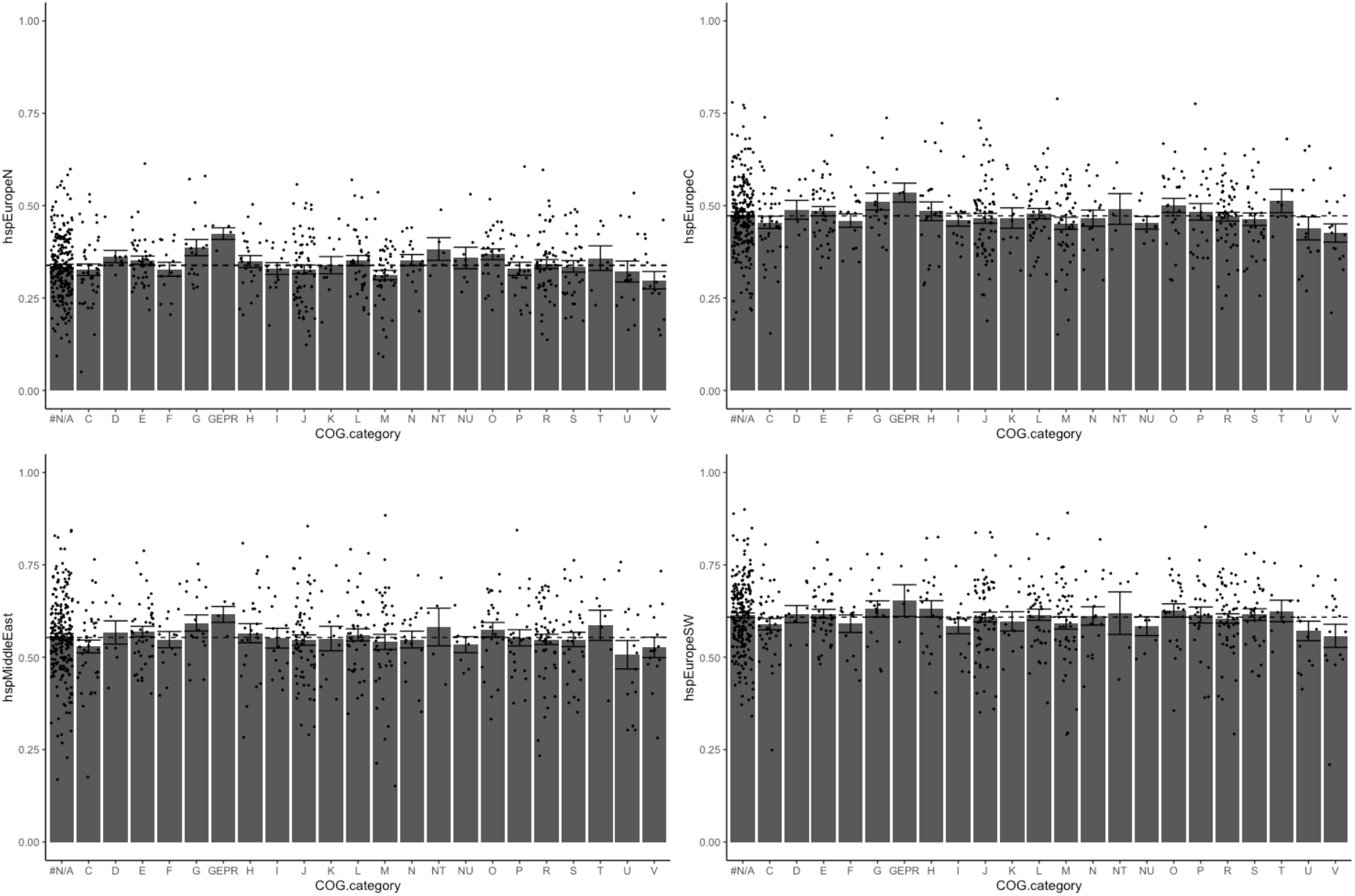
Average African ancestry proportion of genes in particular COG category. Calculated separately for each hpEurope subpopulation. Whiskers show standard error of the average for each category.

**Figure S11.**
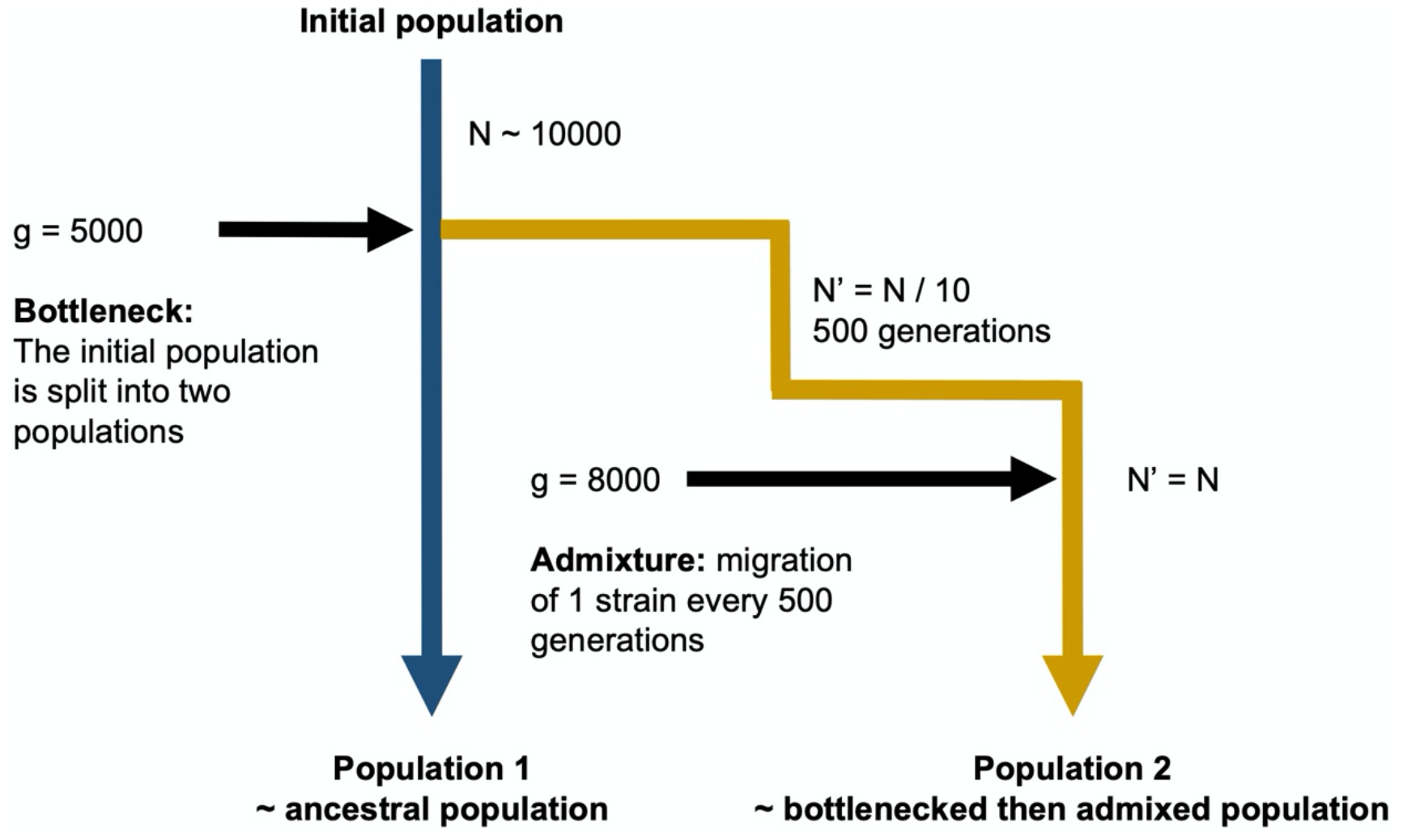
The different steps of the simulation process. Parameters values: N = 10000; genome length = 1.6 Mbp; mutation rate = 5×10^−^7 per bp per generation; deleterious mutations: s = (-0.005, -0.002, -0.001, -0.0005, -0.0002, -0.0001) and they represent 50% of the mutations (the other 50% are neutral mutations). Look at different recombination levels: clonal reproduction (import size per generation = 0bp), intermediate recombination levels (import size per generation = 500bp and 5000bp on average) and nearly free recombination (import size per generation = 50000bp on average).

**Figure S12.**
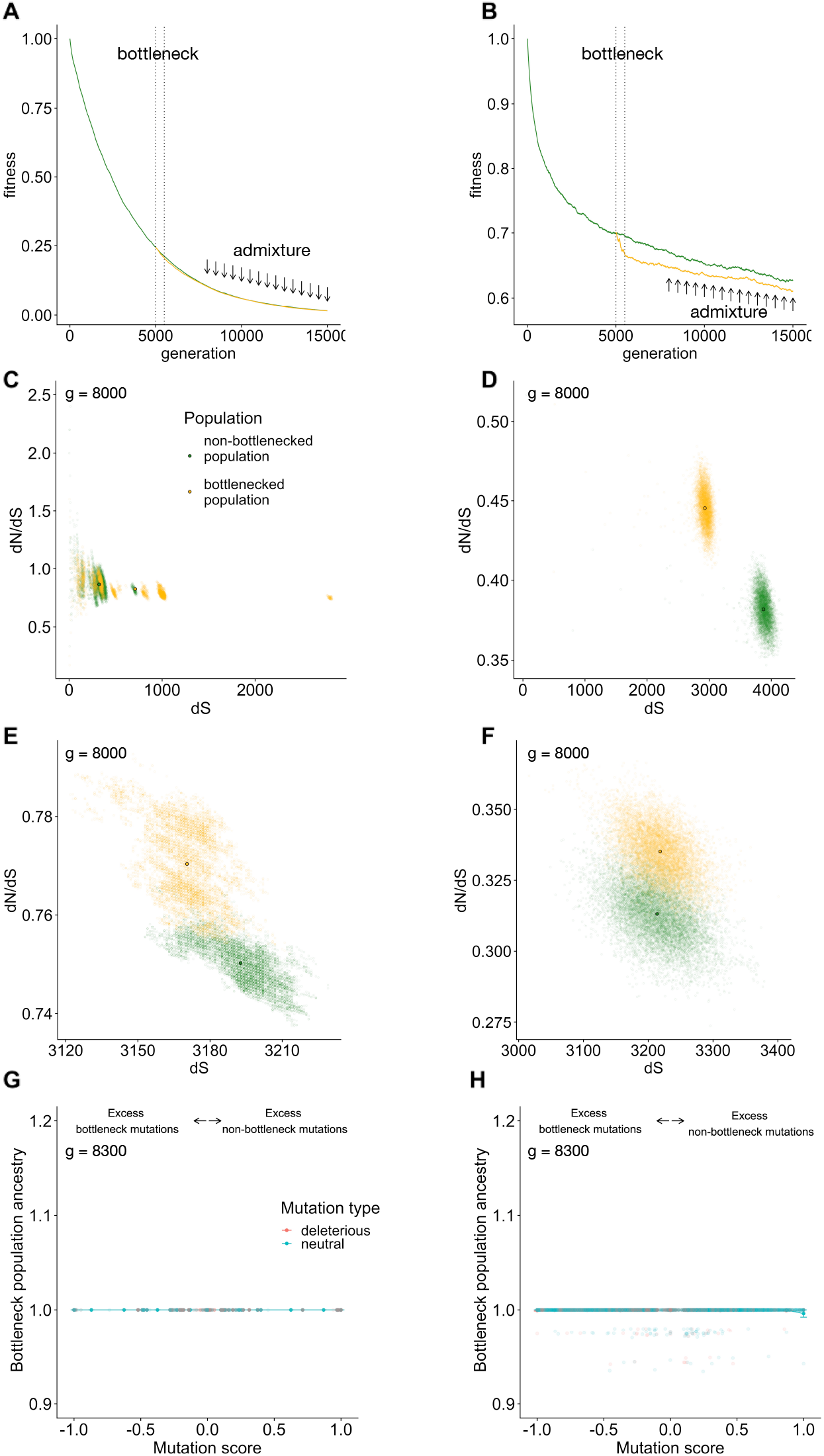
The effect of different recombination rates. (A-B) Average fitness over the generations. (C-D) Within population dN/dS (y axis) plotted against dS (x axis). Semi opaque points show pairwise distances; solid points indicate population means. (E-F) dN/dS calculated to the ancestor plotted against dS for isolates (semi opaque points) and populations (solid points). (G-H) Average bottleneck population ancestry, after the first migration event, plotted against mutation score (frequency in the non-bottleneck population minus the frequency in the bottleneck population before the admixture begins), under no recombination (A,C,E,G) and high recombination (B,D,F,H). The segment represents the bottleneck and the arrow signal the migration events.

